# Imitating the ‘breeder’s eye’: predicting grain yield from measurements of non-yield traits

**DOI:** 10.1101/2023.11.29.568906

**Authors:** Hongyu Jin, Michael C. Tross, Ruijuan Tan, Linsey Newton, Ravi V. Mural, Jinliang Yang, Addie M. Thompson, James C. Schnable

## Abstract

Plant breeding relies on information gathered from field trials to select promising new crop varieties for release to farmers and to develop genomic prediction models that can enhance the efficiency and rate of genetic improvement in future breeding cycles. However, field trials conducted in one environment provide limited insight into how well crop varieties will perform in other environments. As the pace of climate change intensifies, the time lag of developing and deploying new crop varieties indicates that plant breeders will need to make decisions about new crop varieties without knowing the future environments those crop varieties will encounter in farmers’ fields. Therefore, significant improvements in cross-environment prediction of crop performance are essential for creating and maintaining resilient agricultural systems in the latter half of the twenty-first century. To address this challenge, we conducted linked yield trials of 752 public maize genotypes in two distinct environments: Lincoln, Nebraska, and East Lansing, Michigan. Our findings confirmed that genomic predictions of yield can outperform direct yield measurements used to train the genomic prediction model in predicting yield in a second environment. Additionally, we developed and trained another trait-based yield prediction model, which we refer to as the Silicon Breeder’s Eye (SBE). Our results demonstrate that SBE prediction has comparable predictive power to genomic prediction models. SBE prediction has the potential to be applied to a wider range of breeding programs, including those that lack the resources to genotype large populations of individuals, such as programs in the developing world, breeding programs for specialty crops, and public sector programs.

## Introduction

Over the last century, advances in plant breeding combined with improved agronomic practices have been responsible for a six-fold increase in the amount of food that can be produced from a given land area (Grassini, Eskridge, and Cassman 2013; Tester and Langridge 2010). Plant breeders currently employ advanced techniques, including genomic selection to speed the breeding process (Al-Khayri, Jain, and Johnson 2019). However, the start of the plant breeding revolution resulted in no small part, from advances in statistical analyses and a focus on selecting for yield (e.g. crop productivity) in target environments (Breseghello and Coelho 2013; Ceccarelli 2015; Bernardo 2020; Kusmec et al. 2021). These two key factors will continue to be essential for further advancements in agricultural productivity and resilience.

The process of developing new self-pollinated crop varieties propagated from seed can be visualized as an inverted pyramid (Breseghello and Coelho 2013). Thousands or tens of ain, and Johnson 2019). However, the start of the plant breeding revolution resulted in no small part, from advances in statistical analyses and a focus on selecting for yield (e.g. crop productivity) in target environments (Breseghello and Coelho 2013; Ceccarelli 2015; Bernardo 2020; Kusmec et al. 2021). These two key factors will cothousands of new plant genotypes are evaluated at early stages with smaller and smaller numbers advancing to each subsequent stage. Developing new hybrid crop varieties introduces some additional complexities, but they can be set aside for the sake of our initial overview. A plant breeder selects sets of parental crop varieties for crossing and produces large numbers—hundreds to tens of thousands—of true-breeding progeny via multiple generations of self-pollination or in a single generation using a haploid inducer. While the initial number of seeds available to establish a new variety might be dozens or hundreds, these numbers are not sufficient to enable replicated yield plot trials. These initial seeds are propagated in a breeder’s nursery (or less commonly in a greenhouse) to produce sufficient quantities of seed for initial yield trials (Jensen 1989).

However, plant breeders typically produce vastly more new varieties than their programs have the capacity to screen even for early-stage yield trials (Jensen 1989; Xu 2010). Plant breeders will triage a large proportion of their potential new varieties prior to initial yield trials based on simple evaluation metrics. In some cases, the criteria are easily quantified traits (e.g. ear height in maize [*Zea mays*]). In other cases, the criteria might be easy to describe but labor-intensive to collect and not amenable to automation (e.g. flavor in blueberry [*Vaccinium* sect. *Cyanococcus*]) (Gilbert et al. 2015). However, many of these selection steps are based on nebulous criteria or “gut feelings”. These overall evaluation metrics of breeders and experienced farmers were shown to have significant predictive power and can outperform yield measurements during early-stage small plot trials in predicting which lines will perform well in future stages of the plant breeding and selection pipeline.

How can breeders and farmers make predictions from observing a small number of plants of a new variety that outperform observed yield from the same few plants in later large replicated field trials in predicting how the plants will ultimately perform? One model that has been proposed is that skilled practitioners have, over the course of many years, unconsciously learned to detect the correlations between other plant traits, many of which are more stable across genetically identical plants and across environments, as well as crop performance in the farmer’s or breeder’s particular target environment. On a plant-to-plant basis, yield is a relatively variable trait, which is why replicated trials, ideally with each replicate consisting of many dozens to hundreds of plants, are required to accurately estimate the overall genetic potential of a new crop variety. However, many other traits which help determine plant yield are more stable. For example, flowering time measurements collected from one or several plants of a given genotype in a given environment are typically well correlated with flowering time estimates scored from large numbers of plants of that same genotype in the same environment. Hence, a breeder or farmer who has learned the relationships between different stable properties of his or her crop of interest can do a surprisingly good job of predicting which new varieties are worth investing resources into testing them across multiple years and environments, and which varieties might be a waste of resources.

The pool of plant breeders is shrinking. Training a plant breeder requires a substantial investment in resources over many years (Coe et al. 2020). Declines in both government funding for plant breeding research and the number of plant breeding programs have led to reductions in the number of new plant breeders who graduate each year (Stuber and Hancock 2008). In addition, changes in climate and agronomic practices are altering the environments in which future crop varieties will need to be grown. These conditions create the potential for plant breeders to retain or develop less informative mental models for which plant traits best predict good performance in their target environments. As a result, winnowing superior crop varieties by visual inspection may become more error-prone, and there may not be sufficiently trained people available to conduct the work.

Genomic selection acts as an alternative early winnowing method. Here, statistical models trained on genetic and field performance data of previously tested crop varieties evaluate the potential performance of new crop varieties based on the combination of alleles these new varieties carry for the genetic markers used to train the model. Genomic prediction increases the rate of genetic gain in two ways. As an early-stage filter, genomic prediction can discard the bottom tier of lines unlikely to produce a desirable phenotypic extreme, increasing the overall number of individuals that can be evaluated (and hence the overall strength of selection) without requiring a commensurate increase in the number of new lines tested under field conditions. (Belamkar et al. 2018) Thus, genomic selection can act as either a supplement to or replacement for early-stage selection by expert plant breeders (Whittaker, Thompson, and Denham 2000; Meuwissen, Hayes, and Goddard 2001). As a later stage filter, genomic selection can increase the accuracy with which high-performing lines are selected by sharing data on locus effects across related lines within a population. However, due to cost constraints and the logistics of collecting and extracting and sequencing DNA, calling genetic markers, and running predictive models rapidly enough to inform breeding decisions, the routine use of genomic selection at the initial winnowing step is most common in some of the largest private-sector plant breeding programs. Genomic selection is more widely used in both smaller private-sector breeding programs and large public-sector breeding programs later in the downselection process.

Here we tested the feasibility of training machine learning models to replicate the early-stage phenotypic selection process currently performed by plant breeders. We conducted linked trials of roughly 750 publicly available maize genotypes (Mazaheri et al. 2019; Mural et al. 2022) in two distinct environments: Lincoln, Nebraska (NE) and East Lansing, Michigan (MI), collecting grain yield measurements in both environments and hand measurements of several non-yield traits in Lincoln, NE. We confirmed that genomic predictions of yield can outperform the direct measurements of yield used to train the genomic prediction model in predicting yield in a second environment. In addition, we trained another model trait-based yield prediction and which we refer to as the Silicon Breeder’s Eye (SBE). We demonstrate that SBE prediction has comparable predictive power as genomic prediction models. SBE prediction has potential applications to a wider range of breeding programs, including those lacking the capacity to genotype large populations of individuals, such as programs in the developing world, breeding programs for specialty crops, and public sector programs – and to improve cross-environment prediction.

## Results

### Resources that contribute to phenotypic variance

We partitioned the variance in observed grain yield and flowering time (anthesis) from replicated field trials conducted in both locations into five components: G (genetic effect), E (environmental effect), GxE (genotype-by-environment interaction), and residual. We obtained similar results when expressing flowering time in terms of heat accumulation using growing degree days (GDDs) (Figure 1); hence, we used the raw flowering time data in all subsequent analyses. We observed contrasting patterns of variance for flowering time and yield with the environmental component explaining a large proportion of total variance for flowering time (approximately 55% of total variance) but only a small (1.3%) proportion of total variance in grain yield (Figure 1). The GxE interaction played an extremely small role (<2%) in explaining variation in flowering time but explained approximately 13% of the total variance for grain yield across the dataset. In contrast to the small (2.8%) proportion of residual variance explained in the flowering time model, approximately 30% of total variance for yield could not be explained by genotype, environment, GxE, or block effects. This may be due to the reason that the mean difference of the yield between NE and MI was small (∼6%), but the overall yield distributions were different between NE and MI (Kolmogorov-Smirnov *P*=4.9e-6; Figure S5).

**Figure 1.**
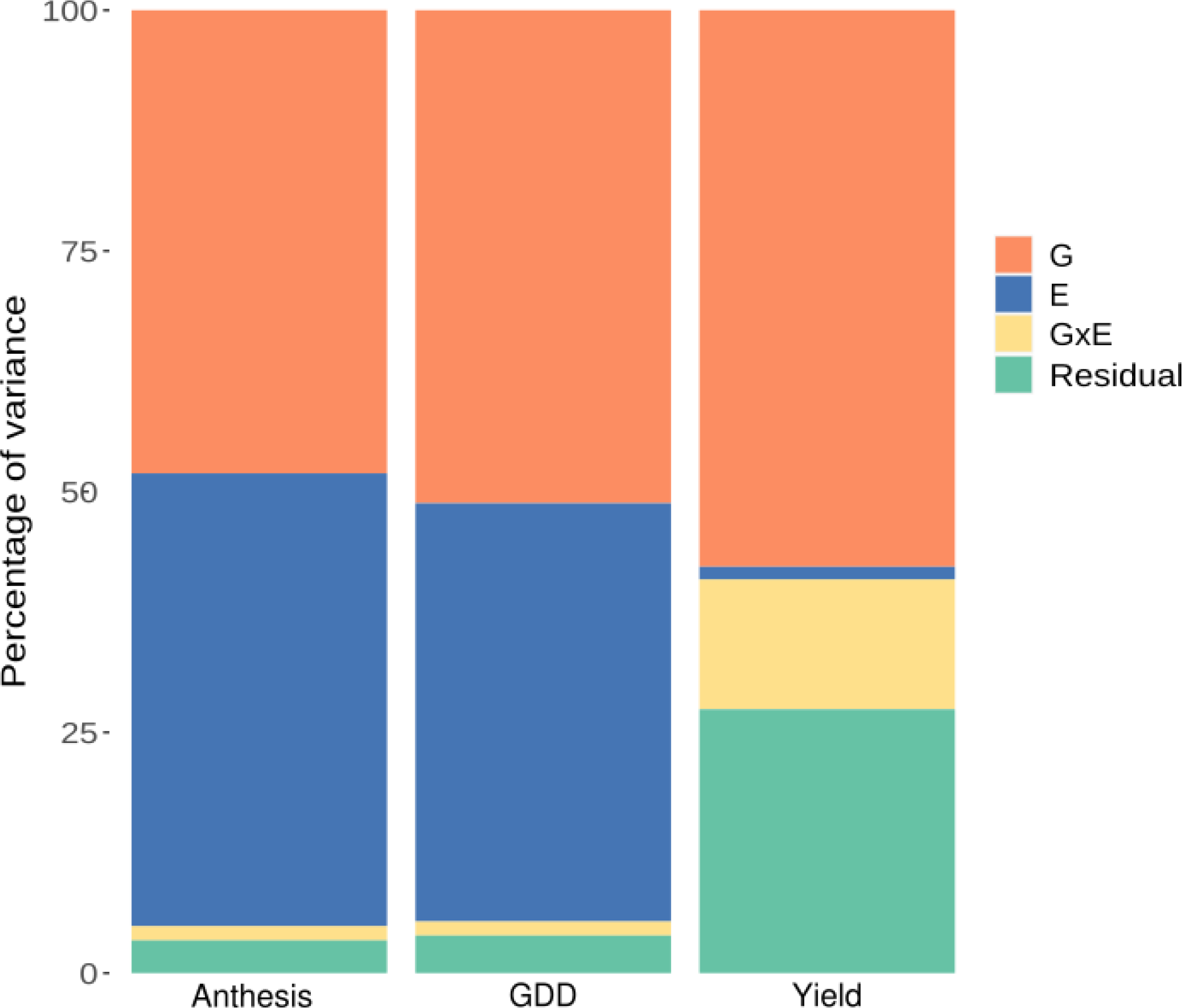
Variance partitioning for flowering time (anthesis), flowering time expressed as growing degree days (GDD) and yield. Colors represent environmental effect (E), genotypic effect (G), genotype-by-environment interaction (GxE), and residual.

Flowering time plays a key role in determining the regional adaptation of maize varieties and productivity across different environments. In the NE field, we obtained the highest yields for genotypes that flowered 70–75 days after planting (Figure S1); in the MI field, we obtained the highest yields for genotypes that flowered 60–65 days after planting (Figure S2). The low degree of GxE interaction for flowering time in this dataset is reflected by the high coefficient of determination (R^2^=0.87; Figure S4) between the observed flowering times in the two environments. The stability of relative flowering time across environments in this dataset suggests that, to the extent GxE in yield can be explained by differences in optimal flowering time in different environments, might be feasible to employ flowering times collected from one environment to predict yield in a second environment.

### Cross environmental yield prediction of the tested genotypes

We built and evaluated three models for predicting the performance of observed genotypes in a previously untested environment: a model predicting yield in the new environment simply by using yield in the observed environment (baseline prediction model); a model predicting yield collected from the new environment using yield in the observed environment combined with genetic marker data (genomic prediction model); and a model predicting yield in the new environment by using yield collected from the observed environment combined with data on other phenotypes collected in the observed environment (SBE prediction model). To perform benchmarking, we first conducted a cross-environment yield prediction using only observed yield data (see Materials and Methods: equation 1). The baseline model simply predicts that yields in new environments will be identical to yields in the observed environment. It is possible to improve the performance of the baseline model simply by generating additional observations of each genotype in the same environment. When only a single observation per genotype in NE was employed to predict grain yield in MI, the correlation between predicted and observed values ranged from R^2^ = 0.37 to 0.40. A model incorporating both replicates produced predictions with a coefficient of determination of 0.45, corresponding to a 5% increase in total variance explained and an approximately 12.5% increase in the proportion of variance explained. This increase was statistically significant (*P*<10^−6^; bootstrap/ANOVA). The inclusion of additional replications decreases the influence of unexpected incidents during the growing season that might influence plot grain yield, which can range from flooding or a low-lying position in the field to microvariations in soil nutrient contents to herbivory by raccoons, deer, or other local fauna. By contrast, in single replicate data, these influences are confounded with genotype means, and in datasets with modest numbers of observations per genotype, unexpected incidents altering the conditions in a single plot can still have a substantial influence on the estimated genotype effect for that variety.

More complex prediction models can share data among genotypes to diminish the influence of unexpected incidents on estimated genetic values for individual lines. One widely used approach is genomic prediction, which builds a model of effects for individual genetic markers based on individuals for which both phenotype and genotype data are available and then predicts phenotypic values for individuals for which only genotype information is available. However, genomic prediction can also improve the prediction of genetic values and, hence, cross-environment or cross-year predictions even when direct phenotypic observations have been collected. We trained a ridge-regression best linear unbiased prediction (rrBLUP) genomic prediction model using yield data collected in NE and a published set of 428,488 single nucleotide polymorphism (SNP) markers. The predicted yield values generated by the rrBLUP genomic prediction model trained using NE yield data exhibited a modest but statistically significant (*P*<10^−6^; bootstrap) increase in correlation with MI yield data (R^2^=0.51, Figure 2B) relative to the observed NE yield data used to train the model (R^2^=0.45).

**Figure 2.**
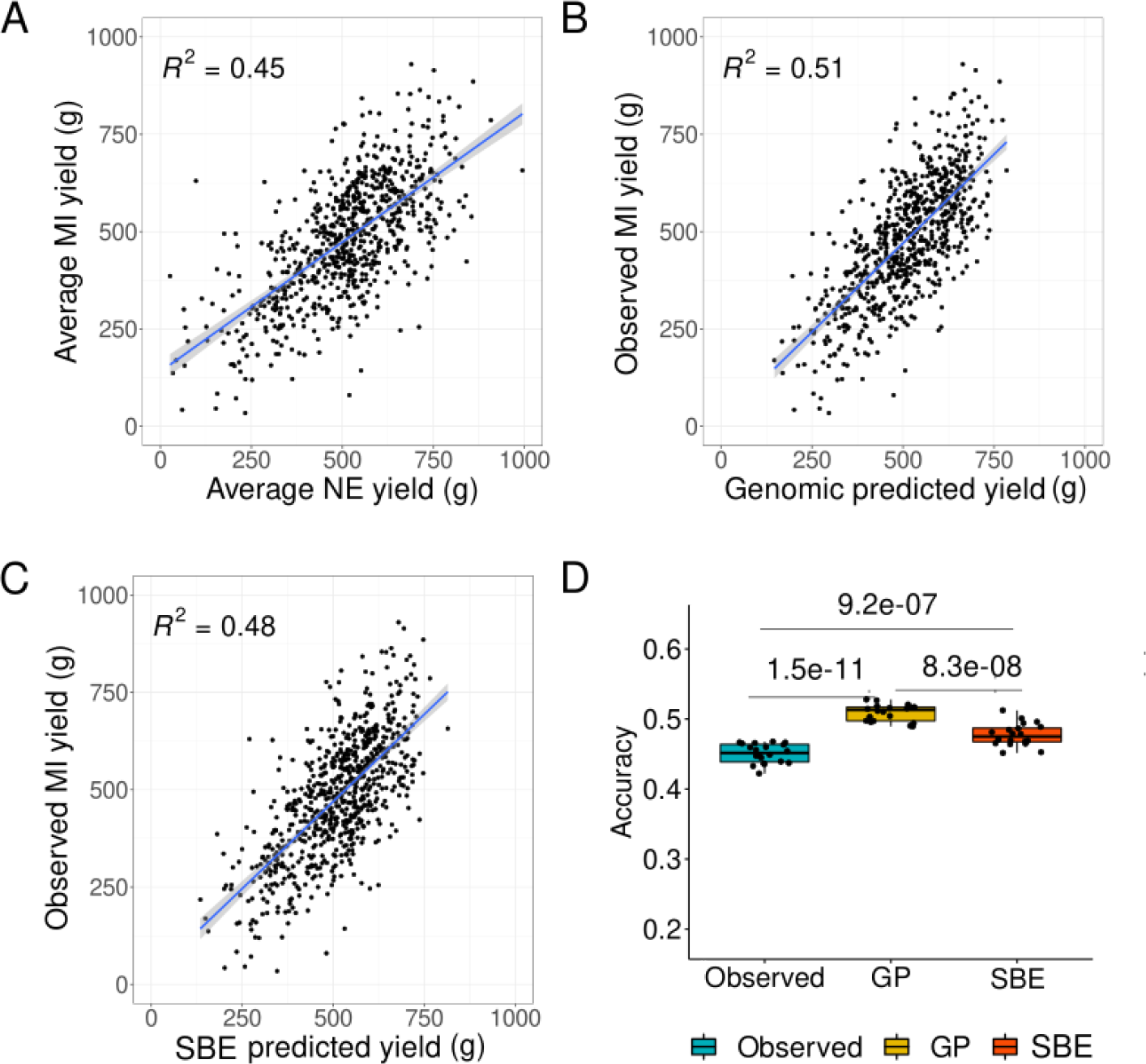
Model accuracy for predicting tested genotypes in untested environments. A, Baseline model using observed data. B, Performance of an rrBLUP genomic prediction model. C, SBE prediction model. D, Bootstrap distribution of prediction accuracies. One-way ANOVA was conducted to compare the means.

The Pearson’s correlation coefficients (r=−0.01 and r=−0.002; Figure S1) between flowering time and yield showed no linear relationship between these two variables. However, in each environment, we observed a substantial nonlinear relationship between these two traits (Figure S1). We calculated a new metric, *x2y*, to quantify the non-linear relationship between the predictors. A number of traits with significant *x2y* relationships with yield can be recorded earlier in the season, before grain yield can be directly scored. In addition, we speculated that, like models built using genetic marker data, models used to predict grain yield based on its relationship to other traits might better predict trait values in new environments by decreasing the influence of unexpected incidents that alter the yield of individual plots. Initially, we attempted to fit an rrBLUP model to predict grain yield using data for 14 phenotypes scored from each plot in NE. This approach performed worse than the baseline model (Figure 2A), with a coefficient of determination R^2^=0.27 (Figure S3). We then employed a random forest model, which may better capture predictive information contained in nonlinear relationships, such as the relationship observed between flowering time and yield (Figure S1). The predictions of a random forest model trained with the same NE grain yield and 14 early-to-mid-season phenotypes described above significantly outperformed (*P*<10^−6^; bootstrap) the baseline model in predicting MI grain yield (R^2^=0.48, Figure 2C), although the improvement in prediction ability using data on 14 other phenotypes to predict grain yield was more modest than that using 428,488 genetic markers to predict grain yield.

The flowering time values associated with the highest grain yields differed between MI and NE (Figure S1). While we observed substantial differences in flowering time between the two environments, flowering times in MI and NE were highly correlated (R^2^=0.87, Figure 6), which is consistent with the small amount of GxE interaction observed for flowering time (Figure 1). As a result, flowering time in NE had a significant predictive value for grain yield in MI (*x2y*=9.93, Figure S1). However, a model trained solely on NE grain yield and flowering time data would learn the wrong relationship to optimally predict MI performance.

### Cross-environment yield prediction of untested genotypes

We evaluated the potential of using plant traits scored in one environment, combined with grain yield scored in a second environment, to predict the grain yield of lines not previously evaluated in the second environment. This scenario frequently occurs in crop improvement and breeding programs, where a much larger set of early-stage lines is evaluated in a single environment, while more advanced lines are replicated in a larger number of environments distributed across multiple remote field sites. The baseline approach for this comparison employed a conventional rrBLUP genomic prediction. The predictions of rrBLUP models trained using grain yield data from 580 maize genotypes in MI and 428,488 genetic markers scored across all 725 genotypes achieved an average R^2^ of 0.45 when the observed grain yields of the 145 maize genotypes used to validate the model were omitted. By contrast, random forest models trained using MI-derived grain yield data for 580 maize genotypes and observed values for 14 early-to-mid-season phenotypes scored in NE achieved an average R^2^ of 0.7, again when omitting observed MI grain yields for 145 lines when training the model. This outcome suggests that detailed phenotyping of large sets of maize lines in a single environment, when combined with data on grain yield for a subset of maize lines in smaller satellite trials across additional environments, can substantially increase the accuracy with which plant performance can be predicted in new environments compared to predicting performance in the same environments based on the same satellite yield results and genomic prediction alone.

### The relationship between intermediate phenotype and the yield

To unmask the contributive intermediate traits to yield, we extracted the feature importance scores from the random forest models and plotted the mean and standard deviation of the scores (Figure 4A). For the random forest models fit to the NE yield data, silking in NE had the highest average importance score (34.4), followed by plant height (20.8) and tassel branch number (20.1). For the models fit to the MI yield data, the top three traits with the highest average importance scores were tassel branch number, silking time, and anthesis time in NE, with scores of 22.3, 20.4, and 17.2, respectively, while leaf number and plant height also had relatively high scores of 15.6 and 15.4, respectively (Figure 4A and B). We plotted the data for some phenotypic features with relatively high importance scores as a function of the yield in both environments (Figure 4C). All scatterplots showed nonlinear relationships between these traits and yields.

**Figure 3.**
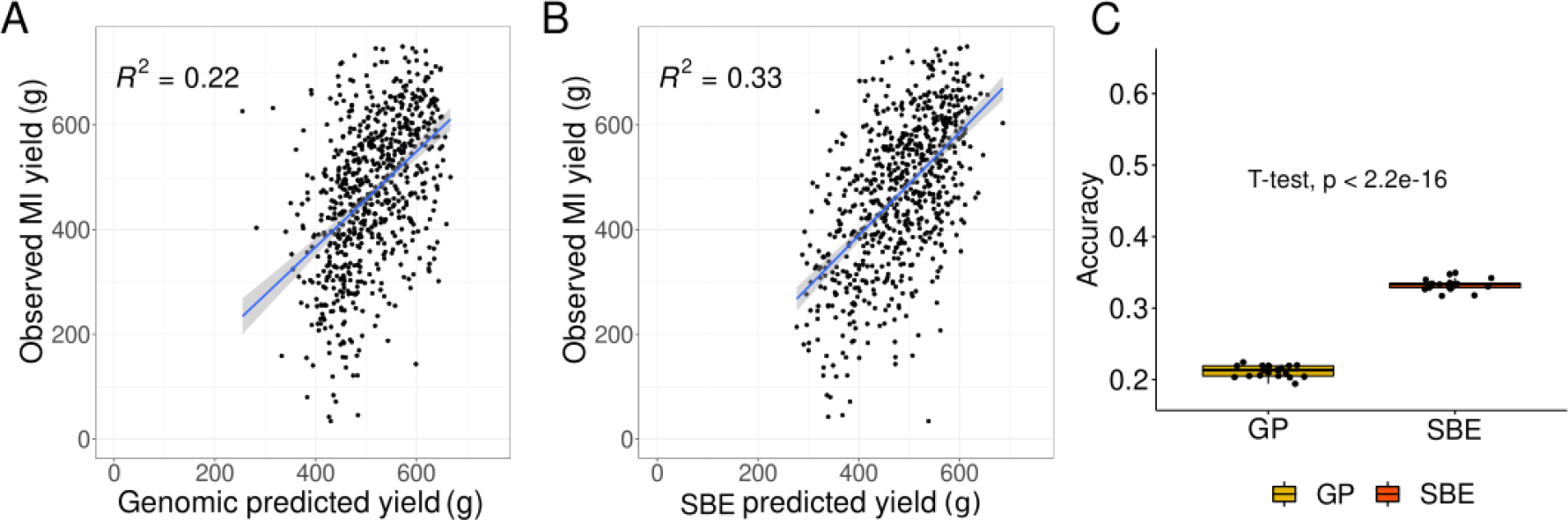
Accuracy of predicting untested genotypes in untested environments or semi-untested environments. A, rrBLUP genomic prediction model. B, SBE prediction model. C, Bootstrap distribution of prediction accuracies. A t-test was conducted to compare their means.

**Figure 4.**
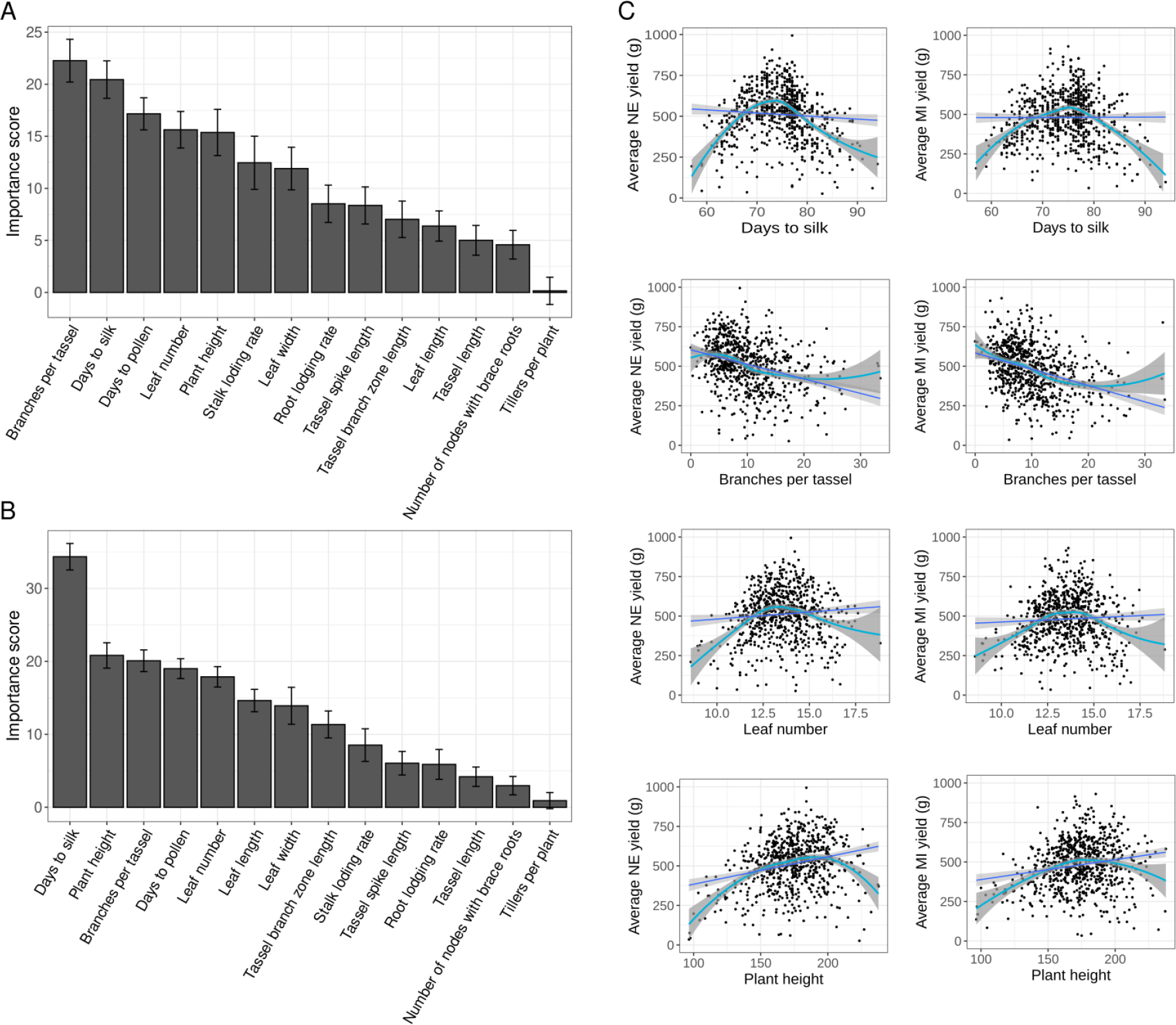
Relationship between intermediate traits and the yield. A, Importance scores of all features in the SBE prediction model predicting NE yield; B, Importance scores of all features in the SBE prediction model predicting MI yield. Important traits are plotted against yield: C, The relationship between NE or MI average yield some of the important traits. Dark blue lines indicate linear regressions and light blue lines indicate polynomial regressions.

## Discussion

Visual selection, which has been practiced for hundreds of years by humans, is still one of the most powerful tools for making selection decisions. We argue that in the absence of experienced human breeders, machine learning algorithms may pattern themselves after human breeders by learning the relationships between phenotypic data and complex traits such as yield. In this study, we tested the accuracies of yield prediction models constructed with genomic or phenotypic data under two prediction scenarios. Both genomic prediction and phenotypic prediction models outperformed the baseline model, which only used the observed data, with statistically significant increases in the accuracy in predicting tested genotypes in new environments. However, when we predicted untested genotypes in new environments, phenotypic prediction integrating the data from other individuals in the new environment yielded higher accuracies than genomic prediction models, whereas the model using only observed data had no prediction power.

The last few decades have witnessed dramatic changes in climate, which are reflected in shifts in temperature as well as increases in volatility (Easterling et al. 2000; Diffenbaugh et al. 2005; O’Gorman 2015). A challenge faced by plant breeders is the need to select varieties for the future, which requires accurate predictions for the performance of new varieties in new environments (Kusmec et al. 2021). Although several models have been developed to incorporate environmental data into predictions (Li et al. 2018; Jarquin et al. 2020), for new field trials, the environmental factors that will be available in the future remain uncertain. The generalization of these models may also be a problem due to the variability of the environmental data and the genetic and physiological interaction effects it may cause. Thus, early to mid-season phenotypic data, which are closely related to plant growth and potentially explain some information from plant-environment interactions, are an informative resource for yield prediction.

Random forest regression is a powerful tool for making predictions (Breiman 2001). As random forest regression is a non-parametric method, it does not require the assumptions of normal distribution, non-collinearity of explanatory variables, and linear relationships between response variables and explanatory variables. This flexibility makes this method suitable for modeling yield based on other phenotypic data. In the comparison between ridge regression and random forest regression for phenotypic prediction, we reached a higher prediction accuracy when using the random forest model. We also attempted to build a prediction model using genetic marker data and the random forest model for the sake of maximum comparability, but this approach would have required the allocation of 684 Gb of memory, which was not practical given local computing resources. Implementing rrBLUP genomic prediction using the same marker set only requires several (<20) Gb of memory, highlighting the value of this computational approach.

The interpretability of machine learning models is defined as their ability to generate knowledge about the relationships (learned by the model) between the data. As a highly interpretable machine learning algorithm, the random forest is able to generate feature importance scores (Breiman 2001; Murdoch et al. 2019). In our random forest-based SBE prediction, we identified the traits that contributed to yield prediction accuracy. Several early to mid-season traits, such as flowering time, plant height, and tassel branch number, played crucial roles in predicting yield. Importantly, with the rapid development of high-throughput phenotyping technologies, such as unmanned aerial vehicles (UAVs), these traits are becoming more easily accessible. Additional traits could potentially be tested and used to increase accuracy of SBE yield prediction models (Alzadjali et al. 2021; Gage et al. 2017; Oehme et al. 2022).

The advantages of SBE prediction make it an ideal approach to be integrated into smaller breeding programs, for several reasons. First, developing and deploying new crop varieties typically takes 7–10 years. As the rate of climate change accelerates, this lag time means that plant breeders will be forced to make decisions about which new crop varieties to advance without knowledge of the future environments these crop varieties will ultimately experience in farmers’ fields (Kusmec et al. 2021). Second, genotyping plant materials to conduct genomic selection could be unaffordable for small breeding programs, especially at the beginning of the breeding program, when larger breeding populations are normally used. Third, experienced breeders might not be available for small or new breeding programs aimed at selecting new crop traits (such as waxy corns) or new crop species (Stuber and Hancock 2008; Coe et al. 2020). With the highly interpretable random forest models used in SBE prediction, new plant breeders will be able to obtain information about which phenotypes to select.

In this study, we demonstrated that SBE prediction, while maintaining similar prediction accuracy, can represent a less expensive substitute for genomic prediction to predict plant performance in new environments. However, the widespread use of hybrids in plant breeding and production makes it inevitable to consider the effects of hybrids on crop yield prediction and selection processes. Given that elite hybrids are more phenotypically similar to one another, more studies should be conducted to assess the performance of SBE prediction using data from hybrids.

## Materials and Methods

### Plant materials and phenotyping

A subset of 752 maize inbred lines from the Wisconsin Diversity Panel was planted on May 6, 2020, in a randomized complete block design (RCBD) with two replications at the Havelock Farm research facility of the University of Nebraska-Lincoln, located at 40.852 N, 96.616 W. Each block consisted of 840 plots, with replicated checks B97, with each plot comprising two rows containing 20 plants. The rows were 7.5 feet in length with a 30-inch row spacing, and there were 2.5-foot alleyways between sequential plots. Further details of this field design are described in Mural et al., (2022). A comparable subset of 761 maize inbred lines from the Wisconsin Diversity Panel was evaluated/screened using an RCBD at the Michigan State University Agronomy Farm, located at 42.709 N, 84.468 W. The seeds were sown in the field on May 25, 2020, with all plots consisting of two rows that were 10 feet in length, with 3-foot alleyways and 30-inch row spacing.

To evaluate flowering time, anthesis and silking dates were recorded following the Genomes to Fieds phenotyping handbook (G2F initiative, 2022). In brief, anthesis was defined as the date when at least 50% of the plants within a plot had exuded anthers shedding pollen at least 50% of the way down the main tassel spike. Silking was defined as the date when at least 50% of the plants within a plot had any amount of silk exuding from their ear primordia. Grain yield was measured for each plot in the NE field as the total kernel weight of 8 randomly selected plants. In the MI field, grain yield was measured as the total kernel weight of 4 randomly selected plants in a plot. The yield data were adjusted to account for a moisture content of 15% for both fields. To ensure comparable values, the yield data from MI were multiplied by two in all analyses.

Other early-to-mid-season phenotypic traits, including flowering time, leaf traits, tassel traits, ear traits, and stand counts were collected in both fields. When possible, edge plants were excluded from data collection to overcome edge effects. Detailed trait lists and collection methods are described in Mural et al., (2022).

### Variance partitioning

The phenotypic data collected from the field trials were employed to evaluate the relative contributions of different factors to variation for two plant phenotypes: flowering time and grain yield. For a given set of phenotypic trait data for all the accessions grown in both the NE and MI environments, the following model was used to describe the observed phenotypic values:

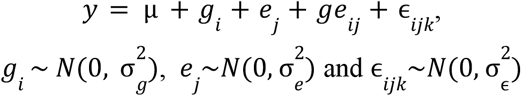

where *y* is the phenotypic value, μ is the grand mean, *g*_*i*_ is the genetic effect of the i^th^ genotype, *e*_*i*_ is the environmental effect of the j^th^ environment, *ge*_*ij*_ is the interaction of the i^th^ genotype and the j^th^ environment, and ϵ is the random error of the i^th^ genotype grown in the k^th^ replicate of the j^th^ environment. The model was implemented in the lme4 package in R (Bates et al. 2014). Then the VarComp() function was used to extract the variance terms and the variances were normalized by the observation numbers. The yield and flowering time (which are strongly affected by the environment) were fitted into the model. Growing degree days (GDD), which integrates flowering time and growing season temperatures, were also calculated for variance partitioning. However, GDD data were not included in the following study due to their highly similar pattern to flowering time data, as shown in Figure 1.

### Genomic and SBE prediction models

We first conducted cross-environment yield prediction without any genomic data or phenotypic data other than yield. For benchmarking, we assumed that maize plants grown in one location would have the same performance in the second location if no additional data were available. In this case, the prediction accuracy was simply calculated as the coefficient of determination R^2^. For genomic prediction, a widely used ridge regression best linear unbiased prediction (rrBLUP) model was employed (Whittaker, Thompson, and Denham 2000), constructed as:

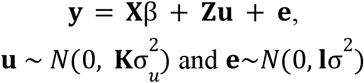

where **y** is a vector of yield phenotypes, **X** is the design matrix (here an identity matrix), β is a vector of fixed intercepts, **Z** is the incidence matrix for the SNP markers, **u** is a vector of random marker effects with covariance matrix 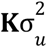, where **K** is the additive relationship matrix from the markers, **e** is a vector of residuals and **I** is an identity matrix with row number equals to **y**. The model was implemented with the *mixed*.*solve* function in the R package rrblup (version 4.6.1) (Endelman 2011).

The SNP markers used for genomic prediction were extracted from an RNA-seq dataset of 942 inbred lines that were genotyped at 899,784 SNPs (Mazaheri et al. 2019). The dataset was then subset to include only the 725 shared lines between the NE and MI fields. SNPs with a minor allele frequency < 0.05 were removed, resulting in 428,488 retained SNPs. A customized script was employed to convert the genotypes into number-coded values, where 0 represents the homozygous reference allele, 1 represents the heterozygous allele, and 2 represents the homozygous alternative allele. A random forest genomic prediction model was trained using the same SNP marker set and the average yield data from NE. The model was implemented in the R package randomForest (version 4.6.14) (Liaw and Wiener 2002) with default parameters. However, due to its high memory usage, this model was not used for the rest of the analysis.

To train SBE prediction models, we used the phenotypic data collected from NE in 2020 (14 early-to-mid-season phenotypic features) as explanatory variables and the yield from either NE or MI as the response variable. The same phenotypic features were also employed as random effects, substituting the marker effects into the rrBLUP model to build a phenotypic prediction model for predicting the yields of tested genotypes in the new environment. However, this model was excluded from downstream analysis due to its inferior performance in prediction.

The *x2y* scores were used to indicate the ability to predict yields based on the early-to-mid-season phenotypes. Briefly, given a set of dependent variable y and independent variable x, the *x2y* score is calculated via three steps: 1) predicting y using a baseline model without x, which in our case was the mean of y; 2) predicting y using a model including x, which in our study was a regression tree; and 3) calculating the percentage reduction of prediction error of step 2) compared to step 1). The scores were calculated using the R package lares 5.1.0 (Lares 2022).

### Prediction scenarios

Two prediction scenarios were conducted and evaluated based on whether the target genotypes or environment were tested. The first scenario involved predicting the yields of tested genotypes in a new environment (MI2020). For this, the genomic prediction model was trained using 80% of the data resampled from all genotypes in the NE data. The prediction was then generated from the same subset of genotypes using the trained model. For SBE prediction, the genetic marker features were replaced by phenotypic features from NE.

The goal of the second scenario was to predict the yields of untested genotypes in a new environment (MI2020). To achieve that, the genomic prediction model was trained using NE data with 5-fold cross-validation. In this approach, 20% of the left-out genotypes or data entries were treated as unobserved testing data, and the coefficient of determination (R^2^) of the model was calculated. Unlike genomic prediction, the 5-fold cross-validation for SBE prediction in the second scenario involved training with NE phenotypic features and MI yield, partially modeling the relationship between the two environments.

Each model was randomized and evaluated 20 times. One-way analysis of variance (ANOVA) was conducted to compare the prediction accuracies between the models in the first scenario, while a t-test was used to compare the models in the second scenario. The code implementing the algorithms is available at https://github.com/JIN-HY/crossenv-yieldprediction.

## Acknowledgements

This project was supported by the U.S. Department of Energy, Grant no. DE-SC0020355 to JCS, DE-SC0023138 to J.Y. and J.C.S., by the National Science Foundation under award IOS-1826781 to J.C.S. and J.Y., by USDA-NIFA under the AI Institute: for Resilient Agriculture Award No. 2021-67021-35329, and Plant Resilience Institute seed funds and Michigan State University startup funds to A.M.T.. The computational resources were supported by the Holland Computing Center of the University of Nebraska, which received support from the Nebraska Research Initiative. We acknowledge Christine Smith, Isabel Sigmon, Kyle Linders and Thomas Hoban for their help in maintaining the NE field. We acknowledge Vladimir Torres-Rodriguez and Nikee Shrestha for informative discussions.

## Author contributions

J.C.S. conceived the experiment. H.J., R.V.M., M.T., R.T., L.N. and A.M.T. conducted the experiment(s). H.J., R.V.M., M.T. and J.C.S. analyzed the results. H.J., R.V.M., J.Y. and J.C.S. prepared the initial draft. All authors reviewed the manuscript.

