## Supplementary figures for "Imitating the ‘breeder’s eye’: predicting grain yield from measurements of non-yield traits"

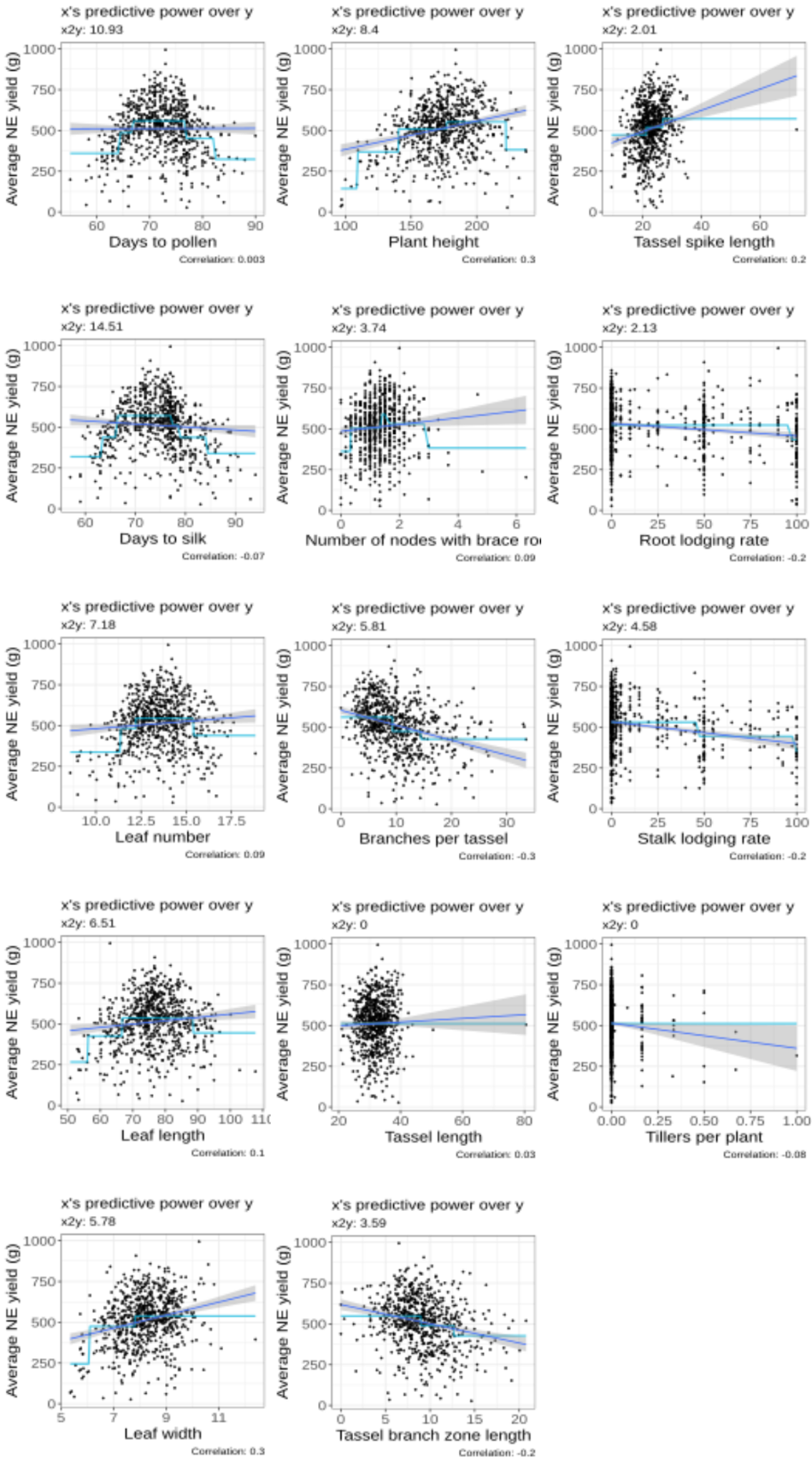

Figure S1. Scatterplots of average yield as a function of phenotypic traits in NE. The darker blue lines indicate the Pearson’s correlation coefficient (r) between two traits, while the lighter blue lines indicate the regression tree models where *x2y* values were calculated.

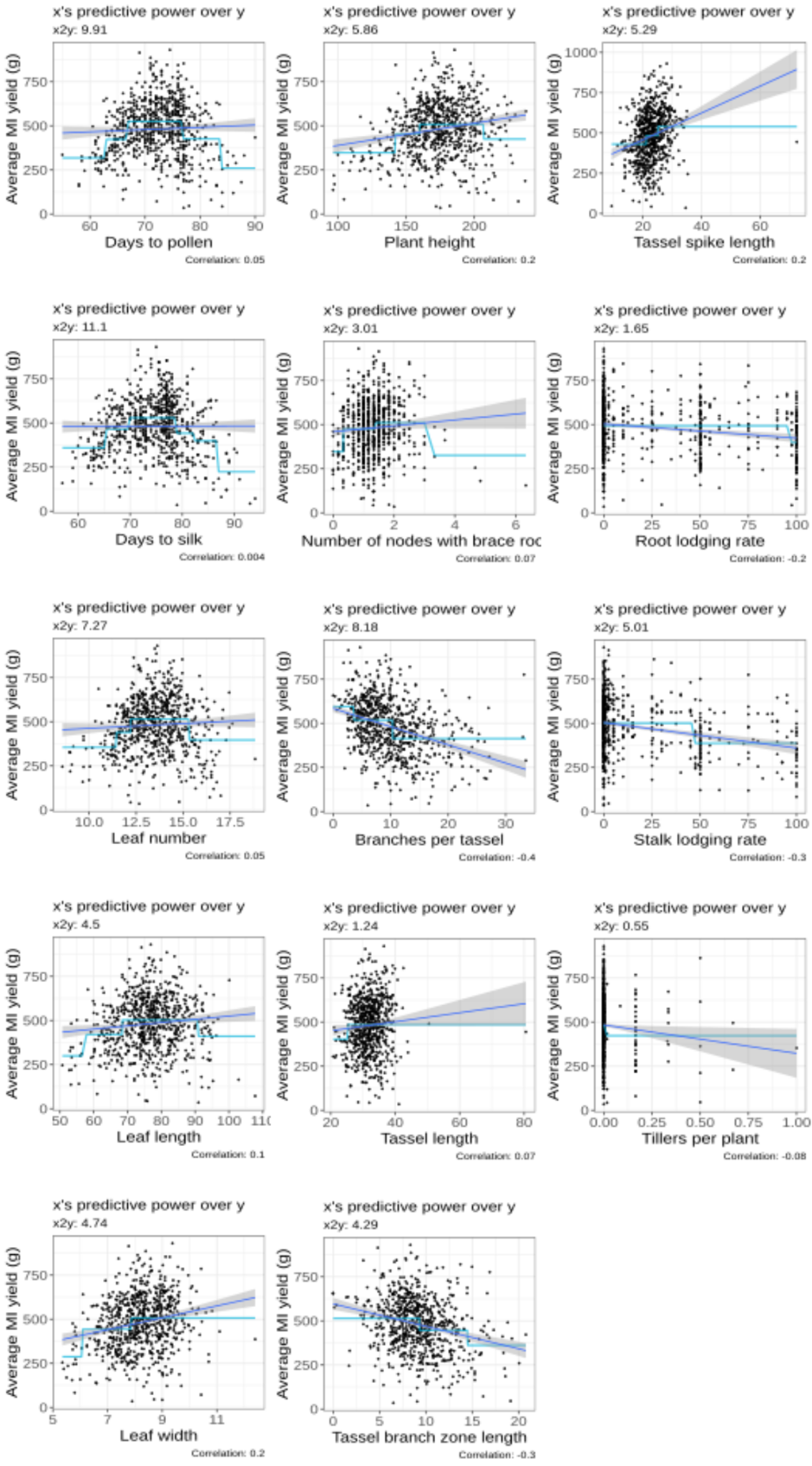

Figure S2. Scatterplots of average yield as a function of phenotypic traits in MI. The darker blue lines indicate the Pearson’s correlation coefficient (r) between two traits, while the lighter blue lines indicate the regression tree models where *x2y* values were calculated.

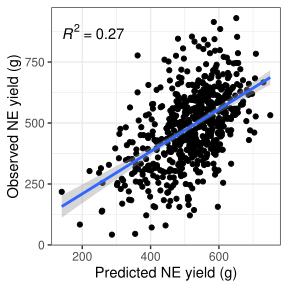

Figure S3. Scatterplot of average MI yield as a function of predicted MI yield generated from an rrBLUP-based SBE prediction model. Predictions were made using tested genotypes.

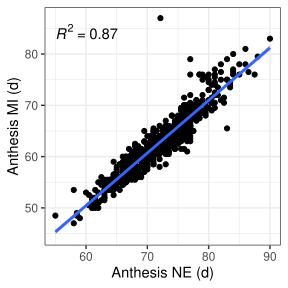

Figure S4. Scatterplot of the correlation between flowering time in NE and MI.

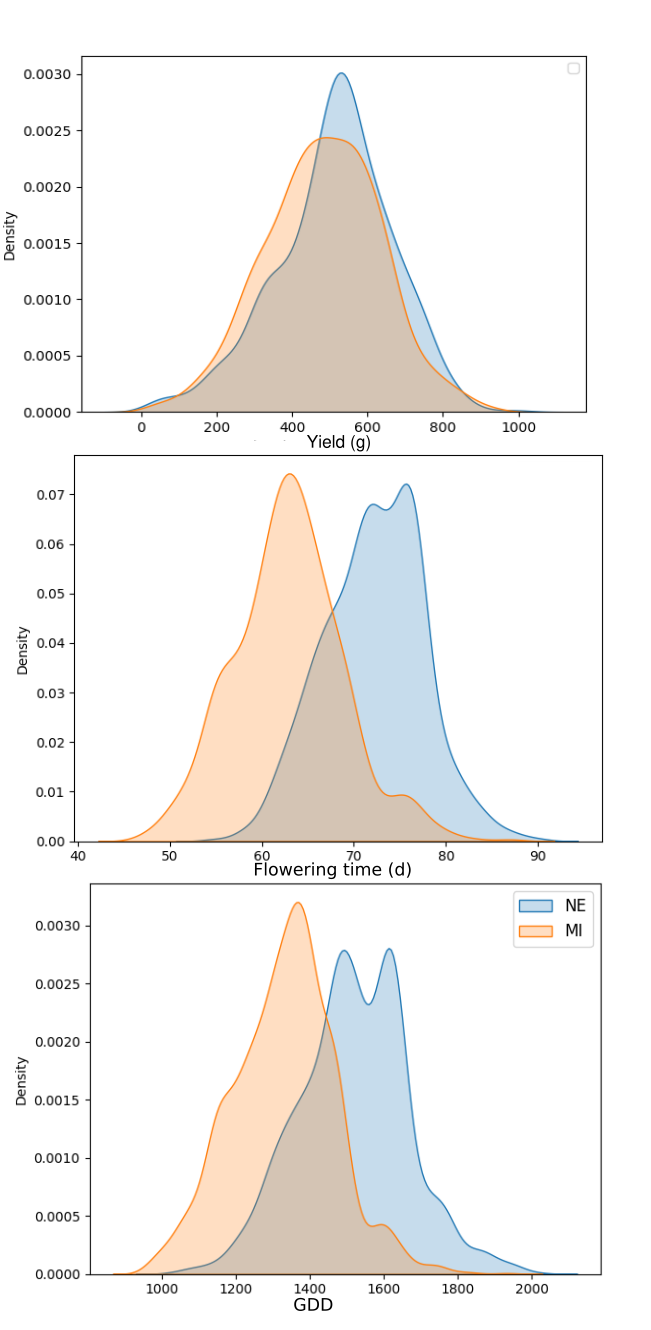

Figure S5. Kernel density plots for yield, flowering time and GDD in NE and MI.
